# Embryonic Atrial Injury Programs a Distinct Substrate for Atrial Fibrillation Characterized by an Anti-Fibrotic Signature

**DOI:** 10.1101/2025.09.16.676656

**Authors:** Fei Xin, Hua Wang, Xinyi Zhao, Xinhui Xu, Jiaheng Cao, Kaiyue Zhang, Yuanyuan Zhang, Jingjing Tian, Yangyang Jia, Changye Sun, Yinming Liang, Xiongwen Chen, Jikui Wang

## Abstract

Atrial myopathy is associated with atrial fibrillation in adults, yet the role of embryonic atrial injury in predisposing individuals to atrial fibrillation remains unclear. Using an inducible, cardiomyocyte-specific diphtheria toxin A-mediated injury mouse model, we found that injured embryonic atria exhibited upregulation of apoptosis and TNF signaling pathways, alongside downregulation of calcium handling and atrial fibrillation-associated genes. Single-cell RNA sequencing revealed two distinct cardiomyocyte populations with differential injury responses. Importantly, although myofibroblasts increased in the fibroblast population, these cells displayed an extracellular matrix gene expression profile distinct from the pro-fibrotic phenotype typically observed in fibrillating adult atria and showed no evidence of fibrosis. Concurrently, monocytes/macrophages were activated and showed upregulated expression of the anti-fibrotic factor *GPNMB*. These findings suggest that adaptive tissue remodeling is counterintuitively prevented from progressing to fibrosis, potentially through GPNMB-mediated mechanisms. This study elucidates molecular pathways linking developmental atrial injury to myopathy and proposes novel strategies to attenuate atrial fibrillation progression.

## Introduction

The atria function primarily as reservoirs, conduits, and contractile chambers. Atrial myopathy is characterized by structural, contractile, electrical, and autonomic remodeling. Various detrimental factors, such as aging, inflammation, oxidative stress, and atrial stretching, can induce atrial myopathy. Progressive atrial myopathy may contribute to atrial fibrillation (AF) in adults (Shen, Arora et al., 2019, Tubeeckx, De Keulenaer et al., 2024).

In the context of aging, structural changes caused by atrial myopathy—including apoptosis, calcium dysregulation, and fibrosis—contribute to the initiation and maintenance of AF. Electrophysiological remodeling is associated with alterations in calcium cycling, ion channels, gap junctions, and autonomic nervous system activity, all of which contribute to the development of a fibrillating atrium (de Groot, Kleber et al., 2025, Shen et al., 2019).

Genetic risk factors increase susceptibility to AF. A significant proportion of patients with early-onset AF carry rare variants associated with myopathy and arrhythmia. Familial AF accounts for more than 5% of all AF patients, indicating a genetic basis for the arrhythmia (Darbar, Herron et al., 2003). Studies demonstrate that genetic mutations lead to familial AF (Chen, Xu et al., 2003, Hodgson-Zingman, Karst et al., 2008, Orr, Arnaout et al., 2016), while genome-wide association studies (GWAS) have identified hundreds of genetic variant loci for AF. Many variants lie near genes implicated in cardiac development or the stress response in adults (Christophersen Rienstra et al., 2017, Nielsen, Thorolfsdottir et al., 2018).

Fetal arrhythmias are a complication in up to 2% of pregnancies (Hornberger & Sahn, 2007). Among these, AF, including familial AF, with fetal onset has been reported (Baldi, Capece et al., 1990, Chao, Ho et al., 1992, Tikanoja, Kirkinen et al., 1998). However, the pathogenesis of atrial myopathy during cardiac development remains largely unknown. Diphtheria toxin-mediated injury of atrial myocytes induces atrial fibrillation, pauses, and complex ventricular ectopy in postnatal mice (Trieu, Mach et al., 2022). Inducible, cardiomyocyte-specific ablation using diphtheria toxin A (DTA) during embryonic development causes embryonic lethality, suggesting severe embryonic cardiac injury (Lee, Morley et al., 1998). To investigate atrial pathogenesis during development, we established an inducible, cardiomyocyte-specific DTA-mediated injury system (iDTA). We found that DTA induction triggers cardiomyocyte apoptosis. At the early injury stage, the top enriched KEGG pathways for upregulated genes involved the apoptotic, p53, and TNF signaling pathways, while downregulated genes were enriched in the calcium signaling pathway. Importantly, *Tbx5* and *Etv1,* genes essential for atrial and cardiac conduction system development, were downregulated in DTA-injured atrial myocytes. Single-cell RNA sequencing (scRNA-seq) identified and characterized two distinct cardiomyocyte populations. Furthermore, we analyzed genes involved in extracellular matrix (ECM) remodeling within fibroblast and myofibroblast subpopulations. We also discovered that monocytes/macrophages (Mono/Mac) were activated in injured iDTA atria, with elevated expression of *Tnf* and *GPNMB* following injury.

## Results

### Tamoxifen-Induced Atrial-Enriched Cre Recombination in Developing Cardiomyocytes

Prior to generating the atrial injury model in developing mouse embryos, we first assessed inducible green fluorescent protein (GFP) expression by crossing *mTmG* with *Myh6-ErCreEr* mice. Membrane-targeted GFP (*mG*) expression in the *mTmG* double-fluorescent Cre reporter mouse is Cre-dependent (Muzumdar, Tasic et al., 2007). The *Myh6-ErCreEr* mouse line induces Cre recombinase activity following tamoxifen (TAM) administration (Sohal, Nghiem et al., 2001).

Pregnant females were intraperitoneally (I.P.) injected with TAM at embryonic days 12.5 (E12.5) and 13.5 (E13.5) on two consecutive days. Embryos were dissected at E14.5 and E16.5, respectively (Figure. 1A). One day post-induction (E14.5), *mTmG*;*Myh6-ErCreEr* embryos appeared grossly normal, exhibiting both mG and tdTomato (mT) fluorescence in the atria and ventricles. Cryosections revealed mG-positive cells within cardiac trabeculae of both atria and ventricles, with stronger signals on the endocardial side (Figure. 1B). Three days post-induction (E16.5), the atria displayed much stronger mG (green fluorescence) signals compared to those at E14.5 (Figure. 1C). In the atria, mG-positive cells accounted for 26.35 ± 3.83% and 43.37 ± 11.44% at E14.5 and E16.5, respectively. In the ventricles, mG-positive cells represented 14.03 ± 12.8% (left ventricle, LV) and 11.01 ± 5.18% (right ventricle, RV) at E14.5; and 0.13 ± 0.21% (LV) and 2.97 ± 2.66% (RV) at E16.5, respectively (Figure. 1D). In contrast, TAM-treated *mTmG* control embryos showed no or very few spurious mG-positive signals in the hearts at all stages (data not shown). These results indicate that the TAM administration protocol induces higher Cre recombination efficiency in the developing atria.

**Figure 1.**
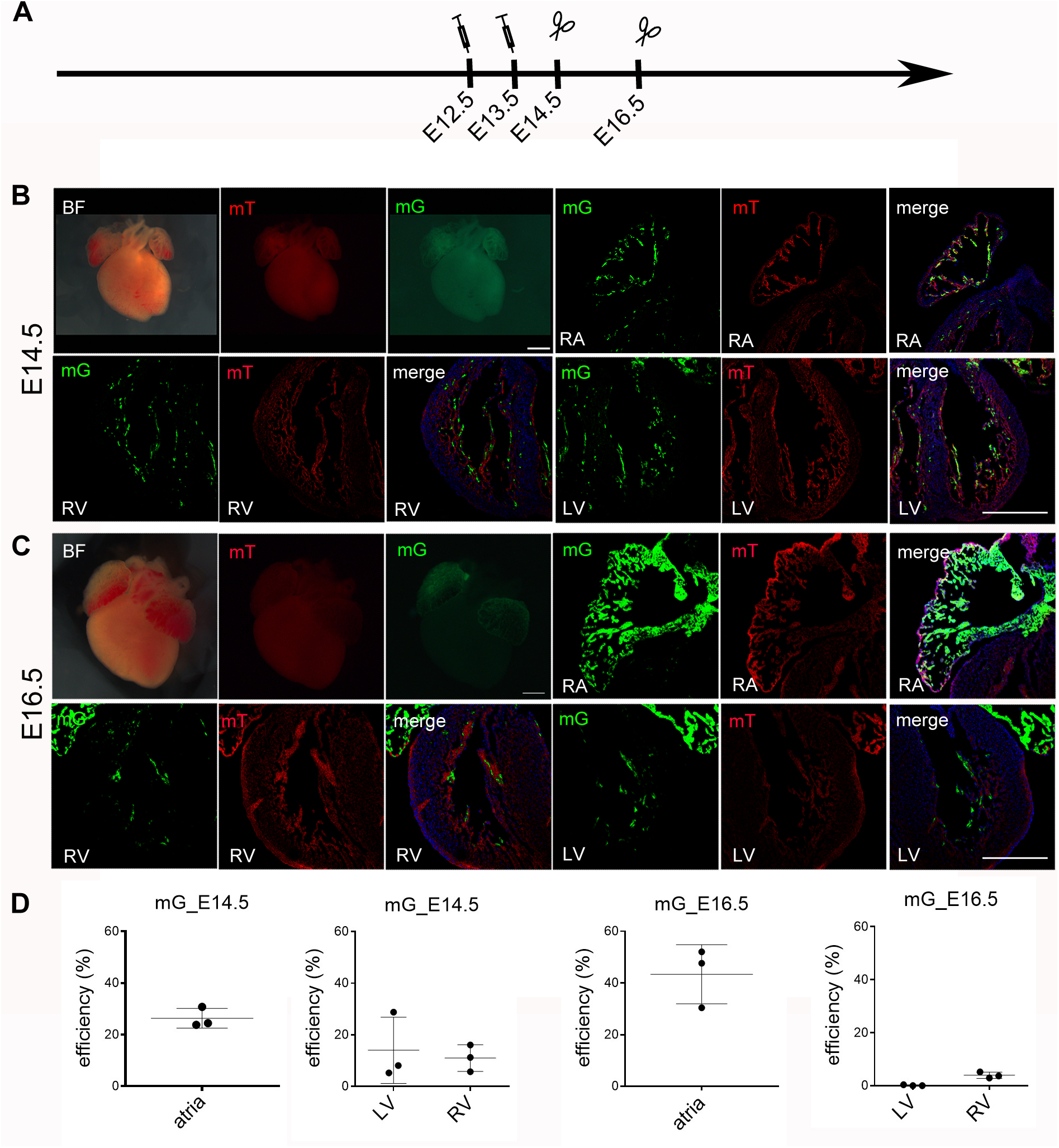
Efficiency of Tamoxifen-Induced Cre Recombination. (**A**) Schematic of the experimental timeline depicting tamoxifen administration and subsequent sample collection to assess Cre recombination efficiency. (**B**) Induced expression of GFP in the heart at E14.5. Scale bars: 500 µm. (**C**) Induced expression of GFP in the heart at E16.5. Scale bars: 500 µm. (**D**) Quantitative analysis of the percentage of GFP-positive cells in the atria and ventricles at E14.5 and E16.5. Data are presented as mean ± SD; n=3. Unpaired Student’s t-test. RA: right atrium; RV: right ventricle; LV: left ventricle.

### DTA-Induced Atrial Myocyte Apoptosis in the Developing Heart

To investigate diphtheria toxin A chain (DTA)-induced atrial myopathy during embryonic development, we established an inducible cardiomyocyte injury model by crossing *Rosa-DTA* mice with *Myh6-ErCreEr* mice. Apoptosis was assessed following TAM administration at E12.5 and E13.5 on two consecutive days.

At E15.5, sporadic TUNEL-positive cells were observed in both *Rosa-DTA* negative controls (Ctrl) and *Rosa-DTA*;*Myh6-ErCreEr* DTA-induced (iDTA) atria. Quantitative analysis revealed no significant difference in the proportion of TUNEL-positive cells (Ctrl vs. iDTA: 0.40 ± 0.20% vs. 1.1 ± 0.98%, *p*=0.3494) (Figure. 2A1-C). Seventy-two hours post-induction (E16.5), clustered TUNEL-positive cells were detected in iDTA atria, with proportions significantly elevated compared to those in Ctrl (Ctrl vs. iDTA: 0.02 ± 0.01% vs. 1.39 ± 0.37%, *p*=0.0114) (Figure. 2D1-F). At E17.5, the percentage of TUNEL-positive cells declined from the levels at E16.5 but remained significantly higher than in Ctrl (Ctrl vs. iDTA: 0.22 ± 0.13% vs. 1.0 ± 0.36%, *p*=0.0220) (Figure. 2G1-I). Collectively, DTA expression efficiently induces apoptosis in atrial myocytes during cardiac development.

**Figure 2.**
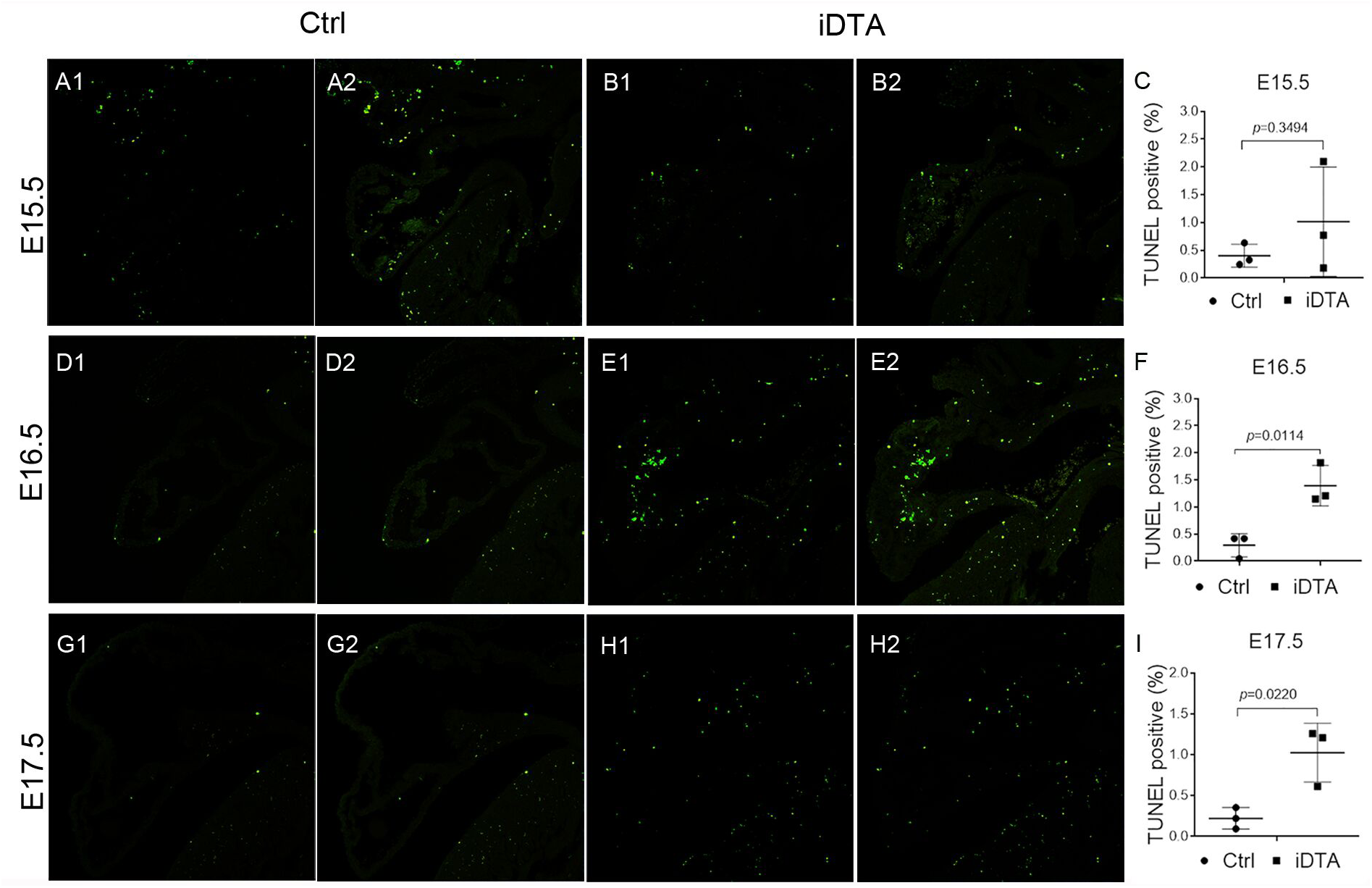
Induced Expression of DTA in Cardiomyocytes Leads to Atrial Apoptosis (**A1-B2**) TUNEL-positive cells shown in the atria of Ctrl and iDTA hearts at E15.5 (Ctrl: A1, A2; iDTA: B1, B2). Scale bar: 200 µm. (**C**) Quantitative analysis and comparison of the percentage of TUNEL-positive cells between Ctrl and iDTA atria at E15.5. Data are presented as means ± SD. Unpaired Student’s t-test; n=3. (**D1-E2**) TUNEL-positive cells shown in the atria of Ctrl and iDTA hearts at E16.5 (Ctrl: D1, D2; iDTA: E1, E2). Scale bar: 200 µm. (**F**) Quantitative analysis and comparison of the percentage of TUNEL-positive cells between Ctrl and iDTA atria at E16.5. Data are presented as means ± SD. Unpaired Student’s t-test; n=3. (**G1-H2**) TUNEL-positive cells shown in the atria of Ctrl and iDTA hearts at E17.5 (Ctrl: G1, G2; iDTA: H1, H2). Scale bar: 200 µm. (**I**) Quantitative analysis and comparison of the percentage of TUNEL-positive cells between Ctrl and iDTA atria at E17.5. Data are presented as mean ± SD; n=3. Unpaired Student’s t-test.

### Bulk RNA-seq Profiling of Damaged Atrial Tissue

To explore molecular alterations in DTA-injured atria, we compared global gene expression profiles between iDTA and Ctrl atria via RNA-seq (E16.5). We identified 648 significantly upregulated genes and 404 downregulated genes in iDTA atria (|log2 fold change| > 0;*p*adj < 0.05) (Figure. 3A, Table EV1). Among the differentially expressed genes (DEGs), 402 upregulated and 159 downregulated genes exhibited fold changes >1.5 (|log2 fold change| > 0.585, corresponding to a 1.5-fold change; *p*adj < 0.05) (Table EV1).

**Figure 3.**
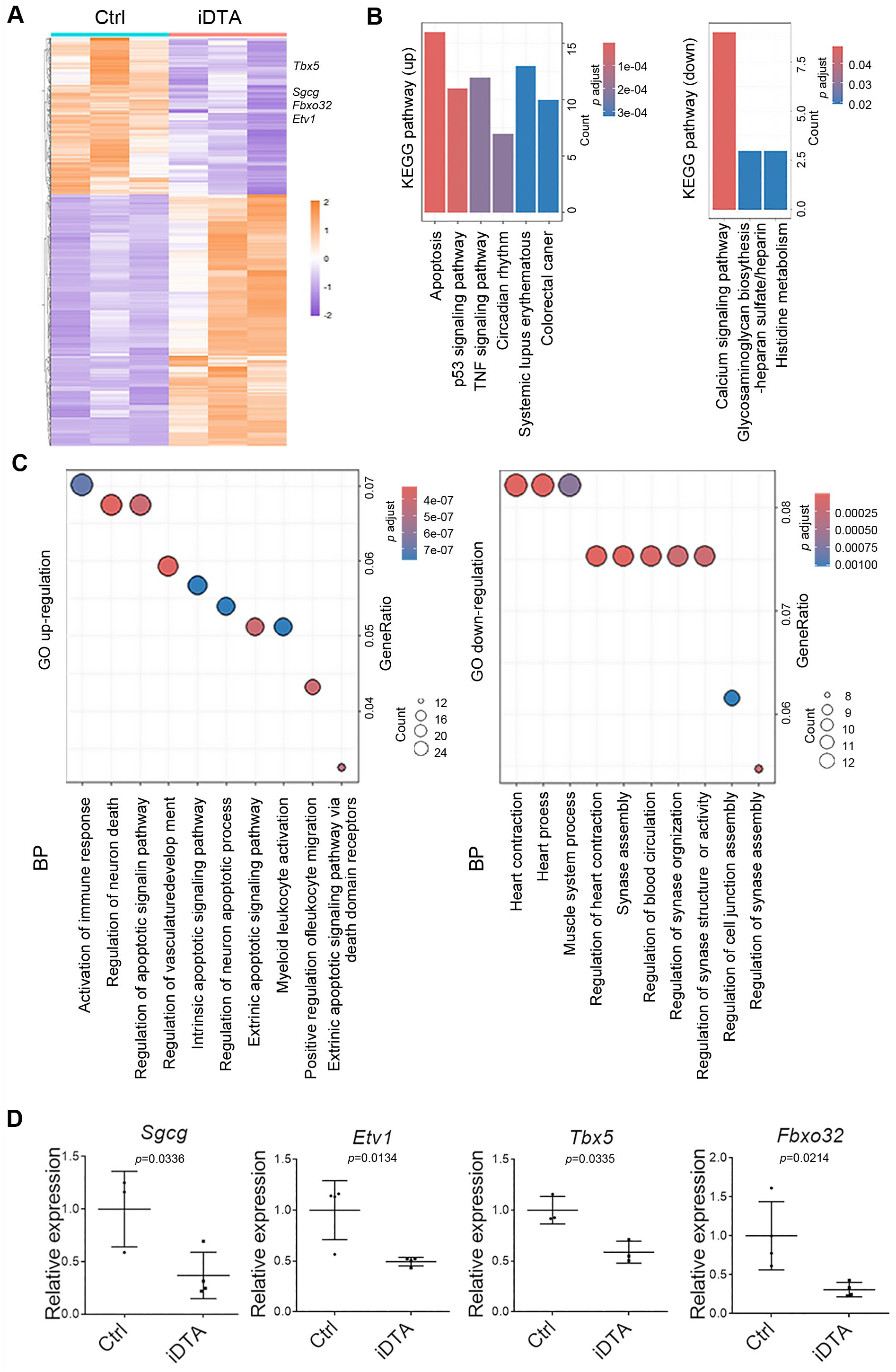
Transcriptional Profiling by Bulk RNA-seq Identifies and Validates Atrial Fibrillation-Associated Genes in Injured Atria (**A**) Heatmap displaying differentially expressed genes in DTA-induced injured atria at E16.5 (648 upregulated and 404 downregulated in iDTA atria; *p*adj < 0.05). (**B**) Enriched top signaling pathways identified by KEGG pathway analysis of upregulated (left) and downregulated (right) DEGs in iDTA vs. Ctrl atria (|log2 fold change| > 0.585, corresponding to a 1.5-fold change; *p*adj < 0.05). (**C**) Top 10 enriched GO Terms associated with upregulated (left) and downregulated (right) DEGs in iDTA vs. Ctrl atria (|log2 fold change| > 0.585, corresponding to a 1.5-fold change; *p*adj < 0.05). (**D**) RT-qPCR validation of AF-associated differentially expressed genes (*Sgcg*, *Etv1*, *Tbx5*, and *Fbxo32*). Data are presented as mean ± SD; n=3-4. Unpaired Student’s t-test.

Kyoto Encyclopedia of Genes and Genomes (KEGG) pathway analysis highlighted upregulation of the apoptosis, p53 signaling pathway, and TNF signaling pathway, alongside downregulation of calcium signaling pathway (Figure. 3B, Table EV2, 3). Gene Ontology (GO) analysis of upregulated DEGs (*p*adj<0.05; |log2 fold change| > 0.585, corresponding to a 1.5-fold change) revealed top enrichment in activation of immune response, regulation of apoptotic signaling pathway, and vasculature development; for downregulated DEGs, predominant enrichment was observed in heart contraction and heart process. (Figure. 3C, Table EV4, 5).

Atrial injury induced by diphtheria toxin has been shown to cause AF in adult mice (Trieu et al., 2022), suggesting potential regulation of AF-associated genes. GO enrichment implicated *Sgcg* in heart process. *Sgcg* encodes gamma-sarcoglycan, a dystrophin-associated transmembrane glycoprotein subunit identified as an AF susceptibility gene by GWAS (Nielsen et al., 2018). A TBX5 mutation cosegregates with familial AF (Postma, van de Meerakker et al., 2008), and atrial *Tbx5* inactivation leads to atrial remodeling and fibrillation in mice (Sweat, Cao et al., 2023). The transcription factor Etv1 is downregulated in patients with reduced ejection fraction; its cardiomyocyte-specific loss induces atrial conduction abnormalities and arrhythmias (Yamaguchi, Xiao et al., 2021). *Fbxo32* encodes a muscle-specific ubiquitin-E3 ligase; its missense mutation results in apoptosis-mediated dilated cardiomyopathy, often associated with cardiac arrhythmias (Al-Yacoub, Colak et al., 2021). DEG analysis revealed downregulation of AF-associated genes (*Sgcg*, *Tbx5*, *Etv1*, and *Fbxo32*) in iDTA atria at E16.5 (Figure. 3A, Table EV1). We validated these findings using RT-qPCR (Figure. 3D).

### Single-Cell Transcriptomic Profiling of Atrial Cell Types

To better understand cellular responses to atrial injury, we generated a single-cell transcriptional atlas of E16.5 atria via scRNA-seq. After quality control, we retained a total of 22,722 cells (Ctrl: 10,628; iDTA: 12,094; average of 2,810 genes/cell). Cell distribution was visualized by Uniform Manifold Approximation and Projection (UMAP) (Figure. 4A). Eleven clusters were identified, ten of which were annotated as distinct cell types using established markers (Figure. 4A, B): fibroblasts, endocardial cells (EndoC), epicardial cells (EpiC), monocytes/macrophages (Mono/Mac), endothelial cells (EC), atrial cardiomyocyte 1 (aCM1), atrial cardiomyocyte 2 (aCM2), smooth muscle cells (SMC), neuron-like cells, and Schwann cells.

**Figure 4.**
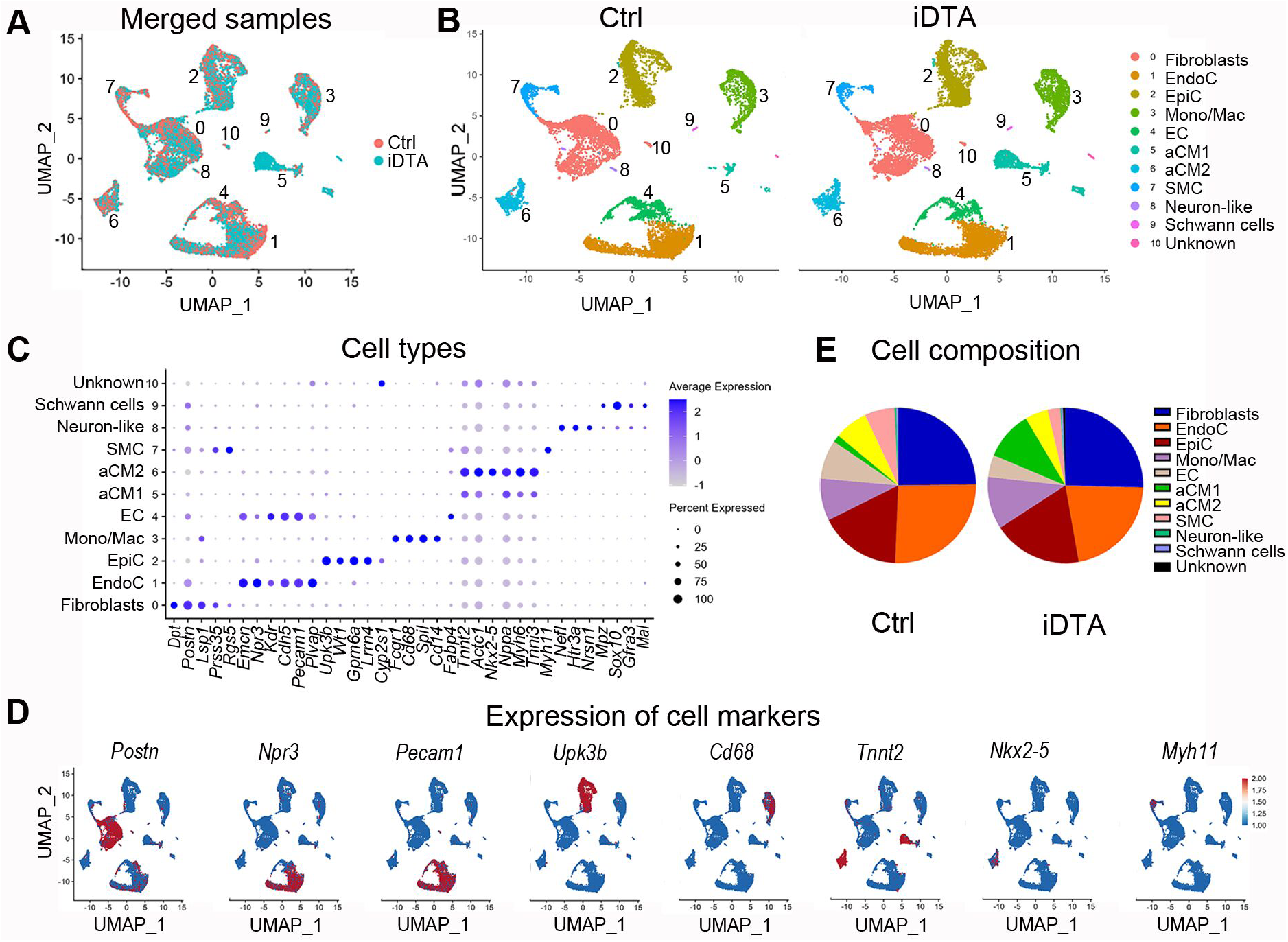
Identification of Cell Types at Single-Cell Resolution in Both the Ctrl and iDTA Atria at E16.5 (**A**) Unsupervised clustering showing the integrated single-cell datasets labelled by the Ctrl and iDTA atria. (**B**) UMAP plots showing the single cells labelled from Ctrl and iDTA atria, respectively. (**C**) Dot plot displaying marker gene expression for cell type assignment. (**D**) UMAP plots displaying the expression of representative marker genes for cell type assignment. (**E**) Pie plots showing the proportions of each cell type in the Ctrl and iDTA atria, respectively.

Fibroblasts were characterized by expression of *Dpt*, *Postn*, *Lsp1,* Prss5 but lacked *Wt1* and *Rgs5* (Buechler, Pradhan et al., 2021, Feng, Bais et al., 2022, Feulner, Wünnemann et al., 2024, McCracken, Dobie et al., 2022, Quijada, Trembley et al., 2021, Xiao, Hill et al., 2018). EndoC expressed *Emcn, Npr3, Cdh5*, *Pecam1*, *Plvap* (Feng et al., 2022, Feulner et al., 2024, Li, Tian et al., 2019, Lu, Wu et al., 2023, McCracken et al., 2022, Rhee, Paik et al., 2021, Zhang, Pu et al., 2016). EpiC specifically expressed *Upk3b*, *Wt1*, *Gpm6a*, *Lrrn4* and *Cyp2s1* (Bochmann, Sarathchandra et al., 2010, Feng et al., 2022, Feulner et al., 2024, Li et al., 2019). Mono/Mac showed high expression of *Fcgr1, Cd68, Spil* and *Cd14* (Feulner et al., 2024, Gula & Ratajska, 2022, Skelly, Squiers et al., 2018, Tondravi, McKercher et al., 1997). EC cells were defined by *Kdr*, *Cdh5*, *Pecam1*, and *Fabp4* (Feulner et al., 2024, Skelly et al., 2018, Zhang et al., 2016). Two atrial cardiomyocyte populations expressed *Tnnt2, Actc1, Nppa, Myh6, and Tnni3*. The aCM2 population exhibited enriched Nkx2-5 and higher expression of *Tnnt2*, Actc1, *Myh6*, *Nppa*, and Tnni3 (DeLaughter, Bick et al., 2016, Feulner et al., 2024, Galdos, Xu et al., 2022, Jia, Preussner et al., 2018, Suzuki, Emoto et al., 2024). SMC highly expressed *Myh11* and *Rgs5* (Amrute, Luo et al., 2024, Feng et al., 2022). Neuron-like cells expressed *Nefl, Htr3a* and *Nrsn1* (Gaetani, Blennow et al., 2019, Ritter & Southard-Smith, 2016, Suzuki, Tohyama et al., 2007). Schwann cells were identified by expression of *Mpz, Sox10, Gfrα3,* and *Mal* (Jessen, Mirsky et al., 2015, Majd, Amin et al., 2023, Widenfalk, Tomac et al., 1998) (Figure. 4C). Representative marker expression for each cell type was visualized in UMAP plots (Figure. 4D).

We next quantified and compared the proportions of these major cell types between Ctrl and iDTA conditions (Figure 4E). In iDTA atria, the proportion of aCM2 was reduced (Ctrl 6.97% vs. iDTA 4.78%), while that of aCM1 was increased (Ctrl 1.51% vs. iDTA 10.19%), suggesting a compensatory response to maintain cardiac development. EpiC, which envelops the heart and migrates inward to differentiate into fibroblasts and smooth muscle cells, was slightly increased (Ctrl 17.04% vs. iDTA 18.59%). The proportion of fibroblasts remained comparable (Ctrl 24.83% vs. iDTA 25.44%). SMC and EC populations decreased markedly (SMC: Ctrl 6.22% vs. iDTA 2.66%; EC: Ctrl 8.10% vs. iDTA 4.56%). EndoC, crucial for chamber morphogenesis via cardiomyocyte interaction (Kim, Nakaoka et al., 2018), showed a substantial reduction (Ctrl 25.78% vs. iDTA 21.80%). The proportion of Mono/Mac increased in iDTA atria (Ctrl 8.78% vs. iDTA 10.93%).

### Characterization of Cardiomyocytes in iDTA Atria

We quantified single-cell expression of AF-associated genes (from bulk RNA-seq) across major atrial cell types (Figure. 5A, Figure. EV1). *Etv1* was expressed not only in atrial myocytes (combined aCM1/aCM2 populations) but also in fibroblasts, EndoC, EpiC, EC, and SMC. We found that the expression of *Etv1* was decreased in atrial myocytes of iDTA embryos. *Tbx5*, *Sgcg* and *Fbxo32* showed specific expression in atrial myocytes, and their expression was downregulated in iDTA atria. We further examined Tbx5 expression at the protein level. Consistent with its transcriptional expression, Tbx5 was found to be decreased in iDTA atria by immunostaining (Figure. 5B).

**Figure 5.**
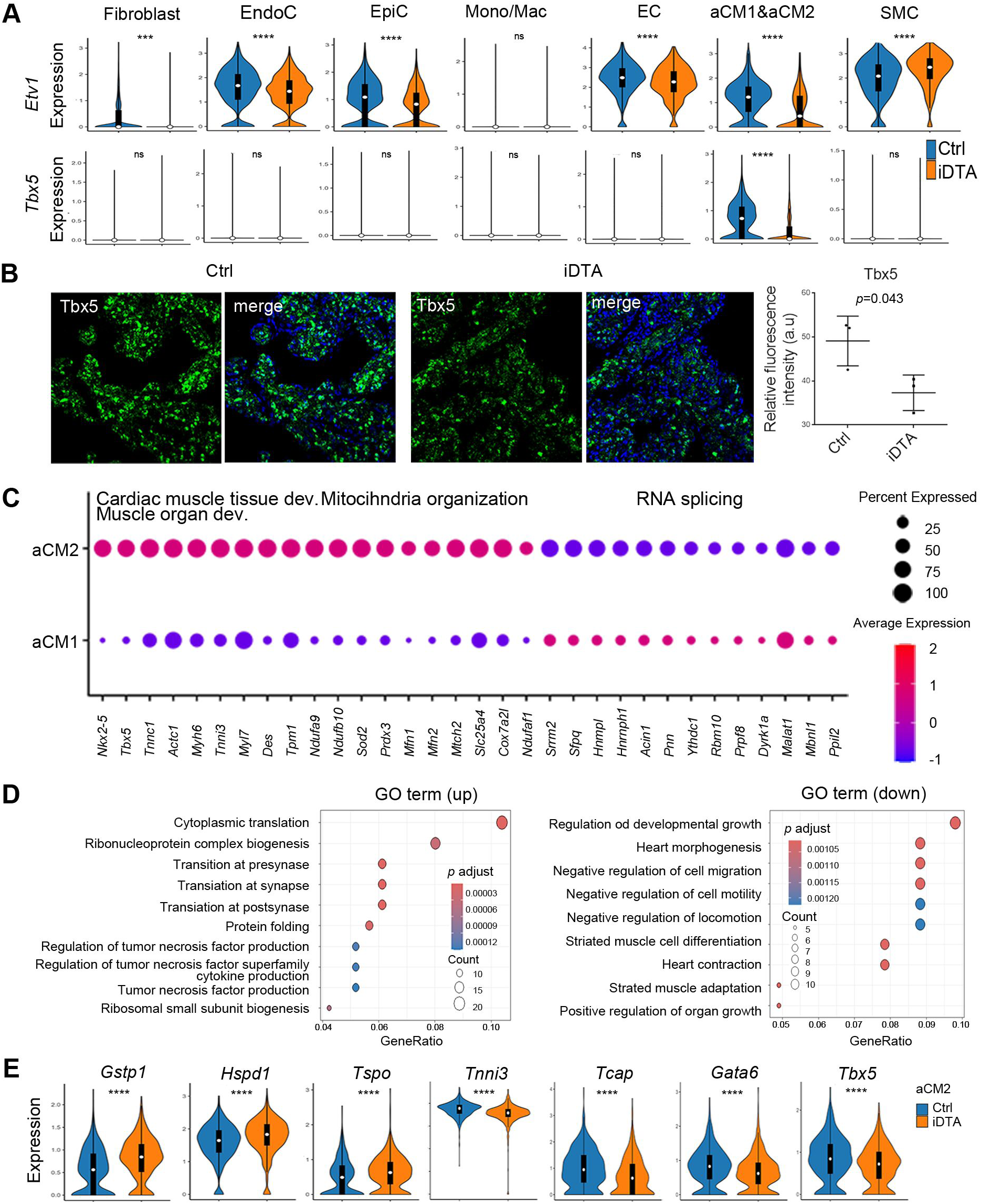
Characterization of Cardiomyocytes following iDTA-Induced Injury (**A**) Violin plots showing the expression of AF-associated genes (*Etv1*, *Tbx5*) in the cardiomyocytes (merged aCM1 and aCM2), as well as in other major cell types. ***: *p*<0.001; ****: *p*<0.0001; ns: no significance. (**B**) Immunostaining images and quantitative analysis of Tbx5 in the Ctrl and iDTA atria. Scale bar: 100 µm. Data are presented as means ± SD; n=3. Unpaired Student’s t-test. (**C**) Dot plot displaying the expression of feature genes from selected top enriched terms associated with DEGs in aCM2 of the Ctrl atria. (**D**) Top significantly enriched GO terms associated with upregulated and downregulated DEGs in aCM2 of the iDTA atria, respectively. (**E**) Violin plots displaying the expression of feature genes from selected top enriched terms associated with DEGs in aCM2 of the iDTA atria. ****: *p*<0.0001.

As unsupervised clustering of scRNA-seq data from E16.5 atria revealed two spatially segregated cardiomyocyte clusters (aCM1 and aCM2), we compared and characterized these populations in Ctrl atria. Differential gene expression analysis showed significantly higher expression of *Nkx2-5* and *Tbx5* in aCM2. Top enriched biological processes (BP) for DEGs downregulated in aCM1 compared to aCM2 included mitochondrial organization, aerobic respiration, and oxidative phosphorylation (Figure. EV1, Table EV 6), suggesting a less mature state for aCM1 (Karbassi, Fenix et al., 2020, Wang, Yao et al., 2020). Moreover, terms related to cardiac muscle tissue development and muscle organ development were also enriched among downregulated DEGs, though not among the top 15 GO terms. Top enriched BP terms for DEGs upregulated in aCM1 compared to aCM2 highlighted RNA splicing in the less mature aCM1 (Figure. EV1, Table EV7). The proportional increase of aCM1 in iDTA atria suggests that the less mature cardiomyocytes may be a potential source for cardiac regeneration in response to cardiomyocyte injury. We next quantified and visualized the relative expression levels of representative genes from these processes across both clusters (Figure. 5C).

Higher *Myh6* expression in aCM2 and the reduced proportion of aCM2 in iDTA atria suggested that aCM2 was likely to be the primary DTA injury target. GO analysis of upregulated genes in iDTA aCM2 revealed top terms including regulation of tumor necrosis factor production and related biological processes (Figure. 5D, Table EV8). The term “regulation of tumor necrosis factor production” encompassed genes involved in detoxification (*Gstp1*), redox stress signaling (*Tspo*), and anti-stress response (*Hspd1*) (Conklin, Guo et al., 2015, Gatliff, East et al., 2017, Sidorik, Kyyamova et al., 2005) (Figure. 5D, E). GO enrichment analysis of downregulated genes in aCM2 from iDTA atria revealed top terms including heart morphogenesis (Figure. 5D, Table EV9). The term “heart morphogenesis” included genes involved in calcium-dependent muscle contraction (*Tnni3*) and sarcomere assembly (*Tcap*), cardiac myocyte differentiation, and conduction system development (*Gata6*, *Tbx5*) (Gharibeh, Yamak et al., 2021, Steimle & Moskowitz, 2017, Zhao, Watt et al., 2008). The relative expression levels of these above-mentioned genes were shown in Figure. 5E.

### Fibroblast and Mono/Mac Response to Cardiomyocyte Injury in iDTA Atria

Cardiac injury induces fibroblast conversion to myofibroblasts—a reparative process for damaged tissue (Younesi, Miller et al., 2024). scRNA-seq analysis detected *Postn* throughout the fibroblast population in both Ctrl and iDTA atria (Xiao et al., 2018) (Figure. 6A). In contrast, *Acta2*, a canonical myofibroblast marker (Johansen, Kasam et al., 2025), was only expressed in a fibroblast subpopulation (Figure. 6B), indicating fibroblast heterogeneity. To characterize the myofibroblast subpopulation, we isolated fibroblasts (cluster 0) from Ctrl and iDTA atria, respectively, and segregated these cells into Acta2-and Acta2+ subgroups. Proportional analysis revealed an increase in the myofibroblast subpopulation in iDTA atria (Ctrl: 74.46% vs. iDTA: 85.08%) (Figure. 6C). Differential gene expression showed that *Tagln* was enriched in myofibroblasts, and its expression in the myofibroblasts of iDTA atria was enhanced compared to that in Ctrl atria (Figure. 6D).

**Figure 6.**
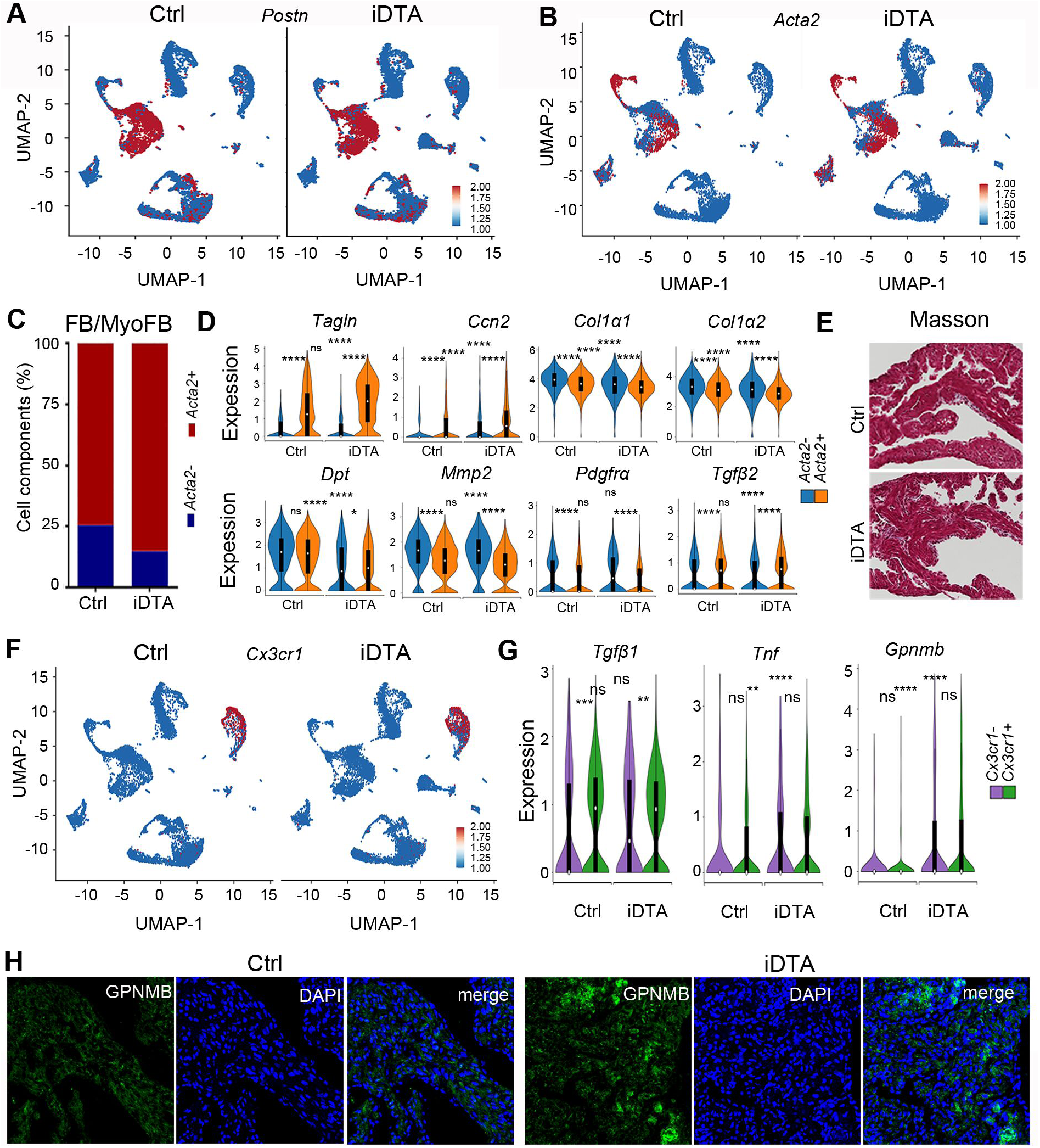
Response of Fibroblasts and Mono/Mac to Injured Cardiomyocytes (**A, B**) UMAP plots displaying the expression of *Postn* in the single cells of the Ctrl and iDTA atria. UMAP plots displaying the expression of Acta2 in the single cells of the Ctrl and iDTA atria. (**C**) Comparison of the proportions of myofibroblasts in fibroblast populations between the Ctrl and iDTA atria. (**D**) Violin plots showing the expression of fibroblast-to-myofibroblast conversion-related and fibrosis-associated genes in the fibroblast and myofibroblast subpopulations of the Ctrl and iDTA atria, respectively. ****: *p*<0.0001; ns: no significance. (**E**) Masson’s Trichrome staining images of the Ctrl and iDTA atria. Scale bar: 100 µm. (**F**) UMAP plot showing the heterogeneity of the Mono/Mac populations identified by the expression of *Cx3cr1* in the Ctrl and iDTA atria. (**G**) Violin plots showing the expression of cardiac repair-associated genes in the *Cx3cr1*-negative and *Cx3cr1*-positive subpopulations of the Ctrl and iDTA atria, respectively. **: p<0.01; ***: *p*<0.001; ****: *p*<0.0001; ns: no significance. (**H**) Immunostaining showing the expression of GPNMB in the Ctrl and iDTA atria. Scale bar: 100 µm.

Atrial fibrosis, a hallmark of atrial cardiomyopathy, is pivotal in AF pathogenesis (Dzeshka, Lip et al., 2015). Fibrosis is characterized by excessive remodeling and accumulation of ECM. In response to cardiac injury, myofibroblasts secrete abundant ECM proteins, which are key components of fibrosis (Stempien-Otero, Kim et al., 2016). *Ccn2*, encoding a matricellular protein essential for fibroblast activation and collagen production (Dorn, Petrosino et al., 2018), was increased in both fibroblasts and myofibroblasts in iDTA atria compared to the corresponding populations in Ctrl (Figure. 6D). *Col1α1*, *Col1α2*, and *Dpt* are ECM genes; their expression was decreased in fibroblasts and myofibroblasts in iDTA atria compared to the corresponding populations in Ctrl atria (Figure. 6D). Expression of *Mmp2*, a matrix metalloproteinase gene, was also decreased in fibroblasts and myofibroblasts in iDTA atria compared to the corresponding populations in Ctrl atria (Figure. 6D). TGFβ stimulates the synthesis of ECM proteins (Frangogiannis, 2022). The expression of *Tgfβ2* was increased in myofibroblasts in Ctrl atria. However, cardiac myocyte injury resulted in only a minimal enhancement of *Tgfβ2* expression in myofibroblasts of iDTA atria compared to that in Ctrl myofibroblasts (Figure. 6D). A recent study demonstrated that inhibition of Pdgfrα signaling reduces myofibroblast expansion at the border of the fibrotic tissue (Protzmann, Zeitelhofer et al., 2025). We found *Pdgfrα* expression was indistinguishable between Ctrl and iDTA atria (Figure. 6D). We further assessed cardiac fibrosis using Masson’s trichrome staining and found no evidence of atrial fibrosis in iDTA atria at E19.5, a later stage of development (Figure. 6E).

Atrial myopathy is closely associated with factors such as atrial inflammation, stretch, and oxidative stress. Clustering analysis identified one Mono/Mac cluster in E16.5 atria (Figure. 4B, C). Gene expression analysis revealed that *Cx3cr1* was expressed in most Mono/Mac cells in both Ctrl and iDTA atria (Figure. 6F). To further characterize the Mono/Mac subpopulation, we isolated these cells (cluster 3) from Ctrl and iDTA atria, respectively, and segregated them into *Cx3cr1*+ and *Cx3cr1*- subgroups. Differential expression analysis showed *Tgfβ1* expression was increased in the *Cx3cr1*+ Mono/Mac subpopulations in both Ctrl and iDTA atria. However, no significant difference was observed in either *Cx3cr1*+ or *Cx3cr1*− subpopulations when comparing Ctrl and iDTA atria. Notably, *Tnf*, an inflammatory mediator gene, was significantly elevated in both *Cx3cr1*+ and *Cx3cr1*- subpopulations of iDTA atria. *GPNMB*, which exhibited low expression in Mono/Mac cells in Ctrl atria, showed increased expression in both *Cx3cr1*+ and *Cx3cr1*- subpopulations in iDTA atria (Figure. 6G). We further performed immunostaining and confirmed the elevated expression of *GPNMB* at the protein level in iDTA atria (Figure. 6H).

## Discussion

In this study, we established an inducible cardiomyocyte-specific injury model using DTA-mediated ablation to explore the pathogenesis of atrial myopathy during cardiac development. Our findings provide new insights into how atrial myocyte injury affects molecular signaling and atrial structure, which may help clarify the potential links to adult AF.

First, our results confirm that TAM-induced Cre recombination efficiently targets developing atria in *Myh6-ErCreEr* mice, with higher recombination efficiency in atria than ventricles. This specificity ensures that DTA-mediated injury primarily affects atrial myocytes, laying a reliable foundation for studying atrial-specific pathogenesis during development.

Using this model, we observed that DTA induction triggers atrial apoptosis in a time-dependent manner: The activation of apoptotic and p53 signaling pathways (from bulk RNA-seq) further supports that DTA injury promotes cell death, consistent with the known role of p53 in mediating apoptosis in response to DNA damage (Phesse, Myant et al., 2014). Conversely, the downregulation of calcium signaling pathways indicates that injury impairs normal calcium handling—an essential process for cardiac contraction —potentially disrupting atrial function during development.

Notably, we found that several AF-associated genes, including *Tbx5, Etv1*, *Sgcg*, *and Fbxo32*, are downregulated in injured atria. *Tbx5* and *Etv1* are not only required for atrial development (Bruneau, Nemer et al., 2001, Shekhar, Lin et al., 2016), but also important for atrial function in the adult heart. *Tbx5* mutation causes abnormal calcium handling and atrial fibrillation, while *Etv1* loss leads to atrial electrical and structural remodeling during pressure overload in mice (Dai, Laforest et al., 2019, Yamaguchi et al., 2021). Their downregulation in iDTA atria suggests that embryonic atrial injury may disrupt the expression of genes critical for atrial structure and electrical function, potentially creating an aberrant developmental trajectory that increases susceptibility to AF later in life. Less is known about the roles of *Sgcg* and *Fbxo32* in cardiac development, and it would be of interest to investigate and validate their function in the atria during cardiac development in animal models.

ScRNA-seq revealed more nuanced cellular responses to injury. The two atrial cardiomyocyte subpopulations (aCM1 and aCM2) showed distinct characteristics: aCM2 represents a more differentiated state, while aCM1 appears less mature. The enriched term regulation of tumor necrosis factor production from upregulated genes (such as *Gstp1*, *Hspd1*, *Tpso*) in aCM2 of iDTA atria suggests that cell stress and inflammation occur in response to atrial injury. The enriched term regulation of heart morphogenesis from downregulated genes (such as *Tnni3*, *Tcap*, *Gata6*, *Tbx5*) in aCM2 of iDTA atria suggests that DTA may repress cell differentiation and maturation. In Ctrl atria, the aCM1 population contains far fewer cells than the aCM2 population. The expansion of aCM1, a less mature population, in iDTA atria suggests a compensatory regeneration in response to atrial injury and cell loss.

Non-cardiomyocyte populations also responded to injury. Fibroblasts, which play a key role in tissue repair, showed an increase in the myofibroblast subpopulation (*Acta2*-positive) in iDTA atria. Myofibroblasts are responsible for ECM remodeling, and our data show elevated *Ccn2* and altered ECM gene expression (*Col1α1*, *Col1α2*, *Dpt*, *Mmp2*), indicating active tissue remodeling. Although TGFβ1 is a known inducer of myofibroblast activation and *Ccn2* expression (Chen, Lam et al., 2000, Dorn et al., 2018, Petrov, Fagard et al., 2002), our pilot data indicate that *Tgfβ1* expression is negligible in fibroblast/myofibroblast subpopulations in Ctrl and iDTA atria (not shown), and the slightly increased expression of *Tgfβ2* in iDTA myofibroblasts likely contributes little to myofibroblast activation and Ccn2 expression (Figure. 6). Given that DTA-induced atrial cardiomyocyte apoptosis—evident as our TUNEL data demonstrated—likely disrupts local tissue integrity and mechanical homeostasis, we reason that such mechanical tension alterations may be the primary driver of fibroblast-to-myofibroblast differentiation. In contrast to the accumulation of ECM and abnormal fibrotic remodeling associated with adult AF pathogenesis, the decreased expression of ECM genes in fibroblast population in iDTA atria suggests that this injury response does not progress to atrial fibrosis during cardiac development, as confirmed by Masson’s trichrome staining.

Mono/Mac cells were activated in injured atria, with elevated *Tnf* and *GPNMB*. Top enriched KEGG of upregulated genes analysis confirmed that the TNF signaling pathway is involved in the pathogenesis of embryonic atrial injury. A Recent study found that *GPNMB* expression is increased in the macrophages of adult infarcted murine heart. Loss of *GPNMB* enhanced the expression of ECM genes (such as *Col1α1*), and worsened fibrosis. While *GPNMB* gain-of-function reduced cardiac fibrosis and enhanced cardiac repair (Ramadoss, Qin et al., 2024). In iDTA atria, the increased Mono/Mac-secreting *GPNMB* most likely downregulate expression and accumulation of ECM genes, repressing fibrosis development.

While GPNMB’s anti-fibrotic role is established in myocardial infarction, its role in AF-associated atrial fibrosis is unknown. Our data suggest that inducing GPNMB expression may mitigate AF-related fibrosis, a hypothesis requiring validation in adult AF models and patients.

Limitations of this study should be noted. First, the iDTA model specifically induces acute injury initiating in cardiomyocytes. This may not fully recapitulate the pathophysiology of human atrial myopathies that originate from or involve other cell types, such as fibroblasts, from the outset. Second, the link between embryonic atrial injury and adult AF remains speculative and will require long-term follow-up studies in animal models.

In summary, our study demonstrates that DTA-induced atrial myocyte injury during development triggers apoptosis, disrupts key signaling pathways, alters cardiomyocyte maturity, activates fibroblast to myofibroblast conversion, and represses fibrosis (likely via immune responses). These findings highlight the vulnerability of the developing atria to injury and identify potential molecular and cellular targets that may contribute to atrial myopathy and AF susceptibility. This study could inform strategies to lessen adult-onset atrial arrhythmia progression.

## Methods

### Ethics statement

All mouse studies were approved by the Institutional Animal Care and Use Committee (IACUC) of Xinxiang Medical University. The mice were housed under pathogen-free conditions in individually ventilated cages, with a 12-hour light/dark cycle and ad libitum access to water and food. All experimental procedures were conducted in accordance with the *Guide for the Care and Use of Laboratory Animals*. Sex was not determined for the experimental embryos.

### Animals

*Rosa-DTA* (JAX: 009669), *Myh6-ErCreEr* (JAX: 005657), and *mT/mG* (JAX: 007576) mice were originally generated as previously described (Muzumdar et al., 2007, Sohal et al., 2001, Voehringer, Liang et al., 2008). Rosa-DTA and Myh6-ErCreEr congenic strains were maintained on a C57BL/6 genetic background, while mT/mG mice were on a hybrid strain of (129X1/SvJ x 129S1/Sv) F1-Kitl+ and CD1. For timed mating, one male and two females were housed together in a single cage. Embryos were obtained from timed matings; the noon of the day when a vaginal plug was detected was defined as embryonic day 0.5 (E0.5). *Rosa-DTA* negative genotypic embryos were assigned as controls, while *Myh6-ErCreEr*; *Rosa-DTA* genotypic embryos were designated as DTA-induced embryos.

### Tamoxifen administration

*Myh6-ErCreEr* males were mated to *Rosa-DTA* or *mTmG* homozygous females. Tamoxifen (Adams-beta) was diluted to 20 mg/mL in corn oil (Sigma-Aldrich) and shaken overnight at 37 °C. Pregnant females at E12.5 and E13.5 received an intraperitoneal injection of tamoxifen at a dose of 200 mg/kg body weight per day. Hearts were then collected in ice-cold PBS at the desired embryonic days for subsequent experiments.

### Lineage tracing

*Myh6-ErCreEr* males were mated to *mTmG* homozygous females. After tamoxifen injection, embryos at E14.5 and E16.5 were dissected in ice-cold PBS. For cryosectioning, embryonic hearts were removed and fixed in ice-cold 4% paraformaldehyde (PFA) for one hour. The hearts were cryoprotected and dehydrated in 18% sucrose for 2 hours with rotation, followed by incubation in 30% sucrose overnight (O/N) at 4 °C. The hearts were then oriented and embedded in OCT compound (Tissue-Tek). Sections (10 µm thick) were cut using a Leica cryostat. The slides were dried for 30 minutes at 37 °C, washed with 1×PBS for 5 minutes to remove OCT, counterstained with DAPI, and mounted with mounting medium. Images were captured using a Nikon Eclipse Ni-U microscope.

### TUNEL assay

Embryonic hearts were fixed in 4% paraformaldehyde (PFA) overnight at 4 °C, dehydrated, and embedded in paraffin. 7µm sections were cut and mounted onto slides. Experiments were performed according to the kit instructions (Promega, DeadEnd ™ Fluorometric TUNEL System). Briefly, after rehydration, slides were treated with 0.85% NaCl to reduce non-specific binding. Tissue permeabilization and digestion were achieved using 0.2% Triton X-100 and 20 µg/mL proteinase K (Roche), respectively. Slides were then incubated with rTdT incubation buffer to label the broken DNA fragments with fluorescein-12-dUTP. The reactions were terminated by immersing the slides in 2X SSC, followed by counterstaining with DAPI. Fluorescent images were captured using a fluorescence microscope.

### Bulk RNA sequencing and data analysis

*Myh6-ErCreEr* males were mated with *Rosa-DTA* homozygous females. DTA expression was induced by administering tamoxifen to pregnant females. Hearts were dissected, and atria and ventricles were separated in ice-cold DEPC-treated PBS at E16.5. Total RNA from atria was isolated from the tissues using TriPURE^TM^ (Roche). Strand-specific libraries were prepared, and RNA sequencing was performed by Novogene Co., LTD (Beijing, China).

Briefly, total RNA quality was assessed using the Agilent 2100 Bioanalyzer, NanoDrop spectrophotometer, and agarose gel electrophoresis. Messenger RNA (mRNA) was isolated from total RNA using poly-T oligo-conjugated magnetic beads. Strand-specific cDNA libraries were prepared using the NEB Next® Ultra Directional RNA Library Prep Kit for Illumina, where the second strand cDNA was synthesized using dUTP instead of dTTP. Paired-end 150 bp sequencing was performed on an Illumina HiSeq X platform.

RNA reads were aligned to the UCSC mouse reference genome (mm10/GRCm38) using HISAT2, and gene counts were calculated using featureCounts. Differential expression analysis was performed with DESeq2. The pheatmap and ggplot2 packages in R were used to generate heatmaps and volcano plots, respectively. KEGG and Gene Ontology (GO) enrichment analyses were conducted using the clusterProfiler R package.

### Single-cell preparation and library construction

Atria were collected from E16.5 embryonic hearts after tamoxifen induction. Fresh tissues were stored in sCelLive™ Tissue Preservation Solution (Singleron) on ice and transported to the company (Singleron Biotechnologies, Nanjing, China) for further processing. Briefly, atria of the same genotype were pooled, minced, and digested with 3 mL of sCelLive™ tissue dissociation solution (Singleron) at 37 °C for 15 minutes. The samples were then mixed with GEXSCOPE® red blood cell lysis buffer (RCLB, Singleron) to eliminate red blood cells. Cell viability was assessed using Trypan Blue.

Barcoding beads were collected from the microwell chip loaded with the single-cell suspension, followed by reverse transcription of mRNA and cDNA amplification. The scRNA-seq libraries were constructed according to the instructions provided in the GEXSCOPE® Single Cell RNA Library Kits. Individual libraries were sequenced on the Illumina NovaSeq 6000 platform with 150 bp paired-end reads.

### Single-cell RNA-seq data processing

Single-cell RNA sequencing data were processed using the Seurat (v5.3.0) and Harmony (v1.1.0) packages in R. Raw gene expression matrices were obtained from Cell Ranger output folders and loaded using the “Read10X” function. For each sample, low-quality cells were filtered by removing those with fewer than 200 detected genes, fewer than 3 cells per gene, or high mitochondrial transcript content (>10%). Following quality control, the resulting Seurat objects were normalized with “NormalizeData”, and highly variable genes were identified using “FindVariableFeatures” with the “vst” method. Data were scaled via “ScaleData”, and principal component analysis (PCA) was performed using the top 15 principal components. Dimensionality reduction was performed using UMAP based on the Harmony-corrected embeddings. For clustering, a shared nearest neighbor (SNN) graph was constructed with “FindNeighbors”, and cell clusters were identified via the Louvain algorithm using “FindClusters”. Cluster identities and group annotations were visualized with “DimPlot”.

Differential expression analysis between experimental groups (Ctrl vs. iDTA) within each identified cluster or specified cell subset was performed using Seurat’s “FindMarkers” function (test.use = “wilcox”), with a log2 fold-change threshold of 0.25. Adjusted p-values were calculated using the Benjamini–Hochberg correction method, and genes with adjusted *p* < 0.05 and absolute log2 fold-change > 0.25 were considered significantly differentially expressed. To further investigate biological relevance, significantly upregulated and downregulated genes were independently subjected to Gene Ontology (GO) enrichment analysis using the “enrichGO” function from the “clusterProfiler” package (v4.10). Bubble plots were generated to visualize the top enriched GO terms, ordered by adjusted p-value and plotted using “ggplot2” with gene ratio and gene count as the x-axis and size aesthetic, respectively.

### RNA isolation and quantitative real-time PCR

Embryonic atria were dissected and total RNA was isolated with RNAiso Plus Kit according to the manufacturer’s instruction. Briefly, complementary DNA was synthesized using RevertAid First Strand cDNA Synthesis Kit according to the manufacturer’s instruction. Quantitative real-time PCR (RT-qPCR) was performed with PowerUp SYBR Green Master Mix Kit according to the manufacturer’s instruction on a Roche LightCycler 96 platform. Relative gene expression levels were calculated using the ΔΔCt method. Target gene expression was normalized to β-actin and expressed relative to Ctrl samples (Table EV10).

### Histology and immunofluorescence (IF)

Embryonic hearts were dissected in cold phosphate buffered saline (PBS) and fixed in cold 4% PFA overnight (O/N) at 4°C. Histology and IF staining were performed on paraffin-embedded 7 µm tissue sections (Table EV10). Briefly, Masson’s Trichrome staining was performed according to the manufacturer’s instruction. GPNMB IF staining was performed following a standard protocol (Table EV10).

IF staining for Tbx5 or Etv1 was performed using Tyramide Signal Amplification (TSA) (Table EV10). In brief, paraffin-embedded sections were deparaffinized and rehydrated. Sections were exposed to 50W LED light for 72 hours at room temperature (RT). To quench endogenous peroxidase activity, sections were incubated with 3% H_2_O_2_ for 30 minutes at RT. Antigen retrieval was performed using sodium citrate buffer. Sections were incubated with primary antibodies O/N at 4°C. After washing, sections were incubated with secondary antibody for 1 hour at RT. Sections were washed again with PBS and incubated with TSA-488 working solution for 10 minutes at RT. After counterstaining with DAPI, samples were imaged on a Leica microscope. Fluorescent intensity was quantified using ImageJ software.

### Statistics

Data are presented as means ± SD. Unpaired t student’s test was used to calculate p values in RT-qPCR and IF experiments. A *p* < 0.05 was considered statistically significant. Exact *p* values are shown where appropriate Statistical analyses were performed on the mean values of Ctrl and iDTA in each group for comparison of differences between two groups. In each experiment, at least 3 samples in each group were used for quantification and statistical analysis. Bar graphs were generated by GraphPad Prism 6.

## Acknowledgements

The authors thank T Chen at National Institute of Biological Sciences for providing mTmG mouse line. This work was supported by the National Natural Science Foundation of China (grant no. 82170237) to J Wang.

## The Paper Explained

### Problem

Atrial fibrillation (AF) is the most common serious heart rhythm disorder. While often associated with aging, it can begin early in life, with fetal-onset cases being reported. However, the atrial pathogenesis of fetal-onset AF remains an unanswered question.

## Results

We developed a novel model of inducible, cardiomyocyte-specific injury in the developing mouse atria. This injury triggered cell death and inflammation, and disrupted genes critical for atrial function (including Tbx5 and Etv1, which are associated with AF). We identified two distinct cardiomyocyte populations with divergent responses to atrial injury. Most intriguingly, we found a paradoxical anti-fibrotic signature: although injury converted fibroblasts to myofibroblasts, they downregulated core ECM genes and failed to produce fibrosis. This was accompanied by activated monocytes/macrophages that secreted higher levels of the anti-fibrotic factor GPNMB, providing a likely mechanism for this protection.

### Impact

Our study suggests that the embryonic heart actively prevents scarring through mechanisms potentially orchestrated by GPNMB. This means early-life injury can create a unique, non-fibrotic substrate for atrial arrhythmia. Our findings point to GPNMB-boosting therapies as a novel strategy to prevent the progressive scarring that drives AF in adults, potentially benefiting a wide range of patients.

## Expanded View Figure legend

**Figure EV1. Characterization of Cardiomyocytes following iDTA-Induced Injury. Related to Figure 5**.

(**A**) Violin plots showing the expression of AF-associated genes (*Sgcg*, *Fbxo32*) in the cardiomyocytes (merged aCM1 and aCM2), as well as in other major cell types. ****: *p*<0.0001; ns: no significance. (**B**) Top significantly enriched GO terms associated with upregulated and downregulated DEGs in aCM1 compared to aCM2 in the Ctrl atria, respectively.

